# Broadly tuned and respiration-independent inhibition in the olfactory bulb of awake mice

**DOI:** 10.1101/002410

**Authors:** Brittany N. Cazakoff, Billy Y. B. Lau, Kerensa L. Crump, Heike S. Demmer, Stephen D. Shea

## Abstract

Olfactory representations are shaped by both brain state and respiration; however, the interaction and circuit substrates of these influences are poorly understood. Granule cells (GCs) in the main olfactory bulb (MOB) are presumed to sculpt activity that reaches the olfactory cortex via inhibition of mitral/tufted cells (MTs). GCs may potentially sparsen ensemble activity by facilitating lateral inhibition among MTs, and/or they may enforce temporally-precise activity locked to breathing. Yet, the selectivity and temporal structure of GC activity during wakefulness are unknown. We recorded GCs in the MOB of anesthetized and awake mice and reveal pronounced state-dependent features of odor coding and temporal patterning. Under anesthesia, GCs exhibit sparse activity and are strongly and synchronously coupled to the respiratory cycle. Upon waking, GCs desynchronize, broaden their odor responses, and typically fire without regard for the respiratory rhythm. Thus during wakefulness, GCs exhibit stronger odor responses with less temporal structure. Based on these observations, we propose that during wakefulness GCs likely predominantly shape MT odor responses through broadened lateral interactions rather than respiratory synchronization.

---

Sensory representations are highly dynamic and can be reformatted to match shifting behavioral objectives^1^. Pervasive inhibitory networks intrinsic to sensory brain regions are well-positioned to broadly regulate temporal patterning and receptive fields, and they are therefore proposed to be critical for state-dependent activity^2–7^. As an example, neural activity in the MOB is state-dependent^8–16^. It has long been suspected that this property is achieved through cortical and neuromodulatory feedback projections accessing the local network of inhibitory interneurons in the MOB^17–20^. Nevertheless, the temporal activity patterns and stimulus selectivity exhibited by these interneurons during wakefulness are not known.

Sensory information is carried to the olfactory cortex via mitral/tufted cells (MTs), which are the principal output neurons of the MOB. Temporal patterning and receptive fields in MTs depend heavily on behavioral state^8–12^. For example, odor representations in awake animals are sparser and more selective than those seen under anesthesia^8, 9^. In the awake state, MTs also exhibit precise bursts of activity that are tightly synchronized to sniffing^11, 12^. Thus, wakefulness in MTs is characterized by smaller, odor-specific neuronal ensembles and strongly respiration-locked output.

The intrinsic MOB network of inhibitory GCs is well positioned to dynamically sculpt these features of MT activity. Granule cells provide feedback inhibition to MTs through a reciprocal dendrodendritic synapse^21–23^. Therefore, it has been widely proposed that feedback inhibition from GCs contributes to temporally patterned activity and stimulus selectivity of MTs^24–27^. Yet, *in vivo* recordings from GCs have been limited to anesthetized animals^16, 28–30^. Two-photon imaging and field potential measurements imply that state-dependent dynamics extend to the GC-MT circuit^31^, but these studies did not resolve the temporal activity and selectivity of individual GCs. Thus, the role of GCs in shaping these features of MT firing is unclear.

We made extracellular recordings from GCs in head-fixed mice while manipulating brain state. The data reveal that under anesthesia odors are represented in GCs by a temporally sparse code dominated by the respiratory rhythm. In contrast, during wakefulness GCs expand their dynamic range and uncouple from the peak of inspiration to fire throughout the respiratory cycle. The increased activity of GCs during wakefulness suggests they have a stronger influence on the sparseness and selectivity of MT receptive fields through broadened lateral interactions. In contrast, the temporal structure of GC firing during wakefulness suggests they have a limited role in respiratory synchronization.

## Results

### Ongoing GC firing characteristics are state-dependent

To measure the state-dependent spiking activity of GCs, we used juxtacellular ‘loose-patch’ methods to achieve extracellular recordings in anesthetized and awake, head-fixed male mice (see Online Methods and Supplementary Fig. 1). Because we were interested in temporal firing patterns with respect to breathing, respiration was also monitored during many experiments (see Online Methods and Supplementary Fig. 2).

Fig. 1a shows data from an example recording in an isoflurane anesthetized mouse, and the subsequently recovered dye fill of the recorded GC. Of the 80 neurons reported here, 50 were definitively identified as GCs by their small soma size and dendritic morphology. The remaining 30 cells were confirmed to be in the granule cell layer but no other morphological data were available. Although there are other cell types found in the granule cell layer^32^, they were very infrequently labeled (2/75 labeled neurons). Moreover, the two data sets were statistically indistinguishable with respect to their basic properties (Supplementary Fig. 3). Therefore, we pooled them for all subsequent analyses.

**Figure 1.**
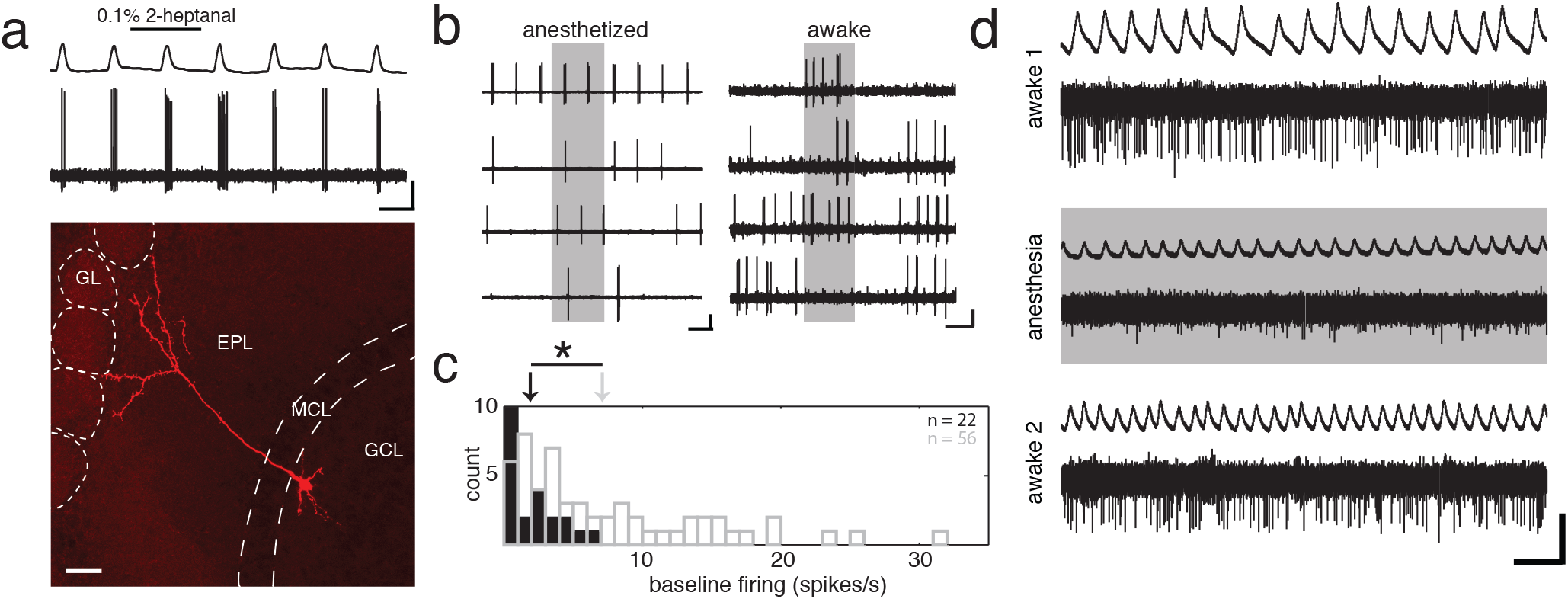
Recording from identified MOB granule cells in the awake and anesthetized mouse. (a) Representative recording from a GC in an anesthetized mouse showing a neuronal spiking trace and respiratory signal above. Scale bar = 2.5 mV/1 s. The lower panel is a photomicrograph of the corresponding neurobiotin filled cell exhibiting hallmark GC features including the apical dendrite extending into the EPL and numerous dendritic spines. Scale bar = 40 µm, GCL = granule cell layer, MCL = mitral cell layer, EPL = external plexiform layer, GL = glomerular layer. (b) Representative traces showing the heterogeneity of odor responses in the awake state. In contrast to anesthetized state GCs, cells in the awake animals variably responded with activation (awake trace 1), delayed onset activation (awake trace 2) and inhibition (awake trace 4) in response to different odors. Scale bar anesthetized = 5 mV/1 s. Scale bar awake = 2 mV/1 s. (c) Distribution of spontaneous firing rates during anesthetized (grey bars, mean = 7.41 ± 7.1 spikes/s) and awake states (black, mean = 1.90 ± 1.9 spikes/s). Spontaneous activity of GCs is significantly increased in awake animals (Mann-Whitney U test, p < 0.01). (d) In select cases, individual cells were recorded during both anesthesia and wakefulness. Spontaneous firing decreases when isoflurane is administered (shaded trace) and returns to pre-anesthesia level when anesthesia is turned off. Scale bar = 1 mV/1 s.

Qualitatively, GCs recorded under anesthesia exhibited low firing rates, robustly rhythmic ongoing activity and odor responses that consisted of modulation of the amplitude of the rhythmic bursts (Fig. 1a). In contrast, GCs recorded during wakefulness exhibited higher levels of aperiodic firing, and responded to stimuli with a mix of increases and decreases in firing rate with variable onset and offset times (Fig. 1b). Mean ongoing firing rates were significantly higher in the awake condition as compared with those seen under anesthesia (awake: 7.41 ± 7.1 spikes/s, n = 58; anesthesia: 1.90 ± 1.9 spikes/s, n = 22; *p* < 0.001, Mann-Whitney U test) (Fig. 1c). These differences appear to reflect state-dependent properties in the same population of neurons rather than properties of distinct populations of neurons. A subset of cells recorded across behavioral state transitions exhibited the same differences between anesthesia and wakefulness (Fig. 1d and Supplementary Fig. 4).

Typically, mice intermittently ran on the freely rotating ball and we measured running velocity during most recording sessions. Although neurophysiological changes to sensory gain and stimulus selectivity have been observed during locomotion^7, 33^, our analysis suggests that the locomotion related firing changes we observed were likely explained by elevated breathing during running (Supplementary Fig. 5).

## During wakefulness GCs exhibit stronger firing rate changes in response to odors

Odor responses in MTs of awake animals are sparser and more selective than those seen under anesthesia. One possible mechanism for this change is greater activity in the GC network facilitating more extensive lateral interactions among MTs. We tested this possibility by comparing the strength and selectivity of odor responses during anesthesia and wakefulness.

Odor responses in awake animals were more robust, broadly tuned and variable in their temporal dynamics than those seen in anesthetized animals. Fig. 2a shows a typical example response histogram for a GC recorded from an anesthetized mouse. Note that tuning curves quantified as mean firing rate changes during (red) and after (green) the stimulus are very flat. A bootstrap procedure testing the significance of individual odor responses (see Online Methods) revealed only a few weak responses. In contrast, robust firing rate changes were commonly observed in GCs of awake mice during the odor. However, qualitative assessment of histograms revealed these robust responses also frequently outlasted the stimulus (Fig. 2b). GC responses in awake mice commonly consisted of a mixture of some excitatory and some suppressive responses to different odors (Fig. 2b). The sign and magnitude of responses during and after the odor were typically not mutually predictive. For awake-state GCs, rank correlation values between tuning curves computed from activity during the stimulus and tuning curves computed from data just after the stimulus were variable but generally low (r = 0.36 ± 0.04, Spearman’s rank correlation coefficient). For most cells (36/49), there was no significant relationship (*p* > 0.05).

**Figure 2.**
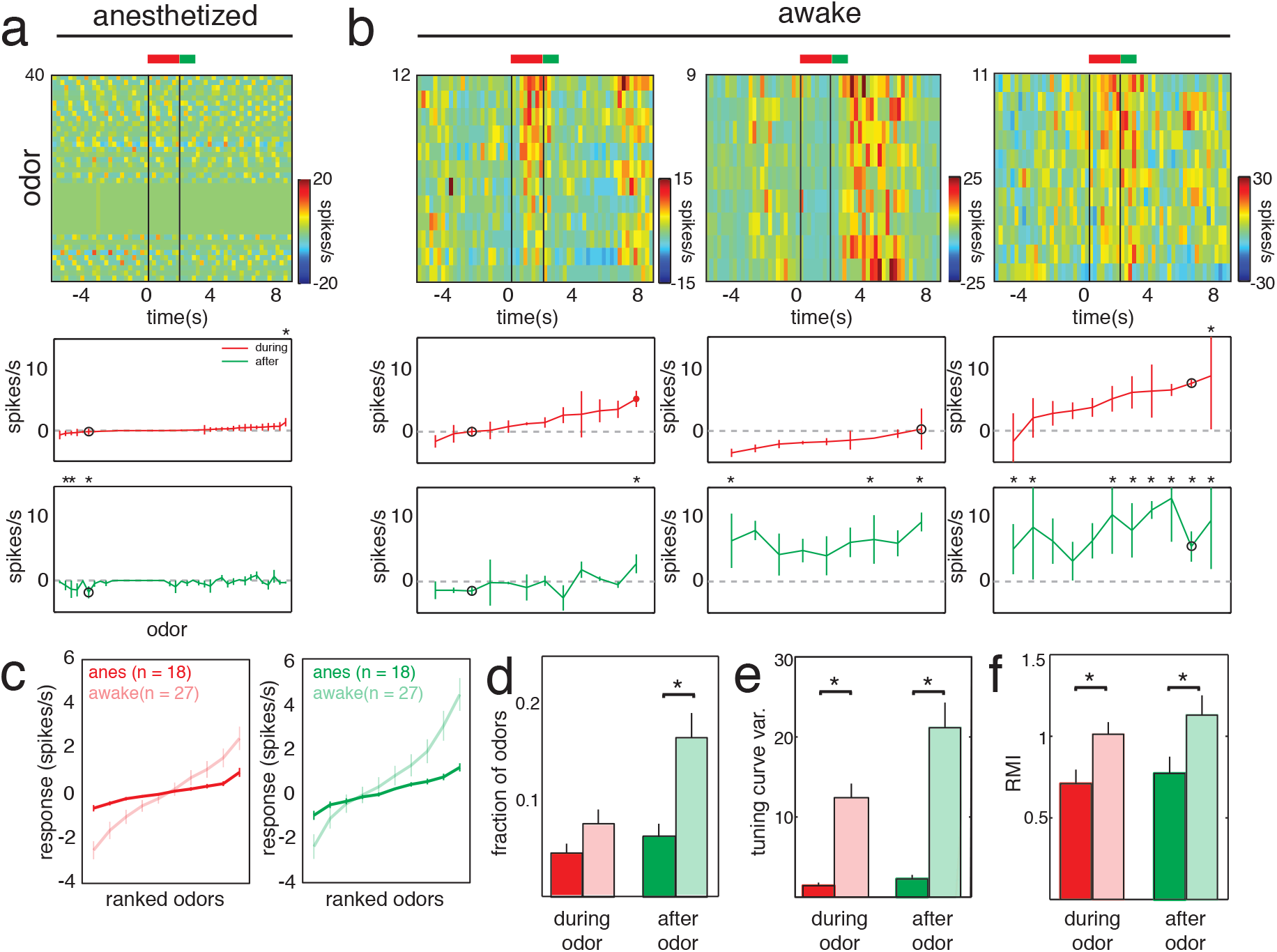
Measures of GC firing rates show stronger and more dynamic odor responses during wakefulness. (a) The top panel is a 2 dimensional peristimulus time histogram (PSTH) with a different odor on every row. Color represents baseline subtracted firing rate binned at 250 ms and rows are sorted with the most excitatory odor response at the top. The vertical black lines denote odor onset at 0 s and odor offset at 2 s. Tuning curves computed from the 2 s during the odor (red) and the 1 s following odor offset (green) are plotted below. Tuning curves are ranked according to strength of the response during the odor, as in the histogram, and they reflect the weak firing rate responses seen under anesthesia. The black circle denoted the response to the blank odor. (b) Representative histograms and tuning curves from three different cells during wakefulness show broader tuning during the awake state. Odors often evoked both excitatory and inhibitory responses both during the odor (left and right curves) and after odor offset (middle and right curves). (c) Ranked mean tuning curves for all cells across 10 standard odors used in both the anesthesia (bold) and awake (light) experiments. Tuning curves calculated from spiking activity during the 2 s odor presentation (red) and for 1 s (green) following odor offset are shown. (d-f) Odor-evoked firing rates were significantly different between anesthesia (bold) and wakefulness (light) comparing three different measures of robustness. The two states showed significant differences (denoted by *, *p* < 0.05, see text for numbers and tests) before and after odor onset in fraction of odors evoking a response, in tuning curve variance across odors, and in response modulation index (RMI). All measures increased in the awake state, indicating that awake state GCs more robustly encoded differences among odors as changes in firing rate.

Mean ranked tuning curves collected for a standard set of 10 odorants were steeper for GCs in awake mice (Fig. 2c) both during the stimulus (anesthesia slope: 0.18 ± 0.1 [spikes/s]/odor, n = 18; awake slope: 0.56 ± 0.3 1 [spikes/s]/odor, n = 27; *p* < 0.001, U test) and after the stimulus (anesthesia slope: 0.24 ± 0.2 [spikes/s]/odor, n = 18; awake slope: 0.76 ± 0.4 [spikes/s]/odor, n = 27; *p* < 0.001, U test). The increase in the overall slope of the tuning curves resulted nearly equally from increased excitation to some odors and increased firing suppression to others (Fig. 2c). Considering all odors presented to all cells, GCs in awake mice responded to a significantly greater fraction of stimuli after odor offset (Fig. 2d) (anesthesia: 6.3 ± 7%, n = 22; awake variance during: 16.8 ± 18 spikes/s, n = 49; *p* < 0.05, U test) and tuning curves collected during both analysis epochs had significantly higher variance (Fig. 2e) (anesthesia variance during: 1.42 ± 1.8 spikes/s, n = 22; awake variance during: 12.2 ± 12 spikes/s, n = 49; *p* < 0.001, U test) (anesthesia variance after: 2.26 ± 2.4 spikes/s, n = 22; awake variance after: 21.1 ± 22 spikes/s, n = 49; *p* < 0.001, U test). This latter analysis does not account for differences in firing statistics between behavioral states. Therefore, we further computed a response modulation index (RMI)^34^ (also see Online Methods) that normalizes the variability across stimuli to the variability across trials separately for each state. Even when correcting for the different levels of noise in this way, the significantly enhanced modulation of firing rate across stimuli in the awake state persisted both during (anesthesia RMI: 0.70 ± 0.4, n = 22; awake RMI: 1.01 ± 0.52, n = 49; *p* < 0.05, ttest) and after the stimulus (anesthesia RMI: 0.77 ± 0.5, n = 22; awake RMI: 1.13 ± 0.85, n = 49; *p* < 0.05, ttest).

These data show that GCs respond more strongly and to a broader range of odors in the awake state as compared to the anesthetized state. Even after correcting for different firing statistics, our analysis reveals that GC spiking in response to odors has a greater dynamic range across stimuli. These data reveal that GCs exhibit stronger, more broadly-tuned responses in the awake as opposed to the anesthetized state. Therefore we conclude that the awake state is characterized by amplified lateral interactions between MTs via GCs. Further, our finding that GC responses frequently continue beyond the end of the stimulus raises the possibility that during wakefulness GCs reflect non-sensory events or cortical influence, as opposed to bottom-up sensory input.

## Respiratory coupling of GCs is state dependent

Under anesthesia, activity in MTs and GCs is strongly coupled to respiration^28, 35–40^. During wakefulness, MT spikes are also locked to breathing and sniffing^11, 12^, however the temporal structure of GC firing in this state is unknown. Inhibition from the GC network during wakefulness may play an important role in enforcing respiratory locking of spikes in MTs. We therefore examined whether GC firing is coupled to breathing, whether it is coupled to a characteristic phase of breathing, and whether this coupling changes with behavioral state.

We quantified the strength of coupling between GC firing and the respiratory cycle using a metric *r* that ranges from 0 to 1, with 0 corresponding to even distribution of spikes throughout the breath cycle and 1 corresponding to all spikes occurring at the same point in the breath cycle^41^. To our surprise, during wakefulness GCs were much more weakly coupled to respiration then they were under anesthesia. Fig. 3a (left panel) shows respiratory phase histograms for representative (near mean) GCs recorded during anesthesia (black) and during wakefulness (gray). GCs recorded during wakefulness were significantly more weakly coupled to the breathing rhythm than GCs recorded under anesthesia (Fig. 3a, right) (anesthesia r: 0.59 ± 0.2, n = 16; awake r: 0.25 ± 0.24, n = 18; *p* < 0.001, U test).

**Figure 3.**
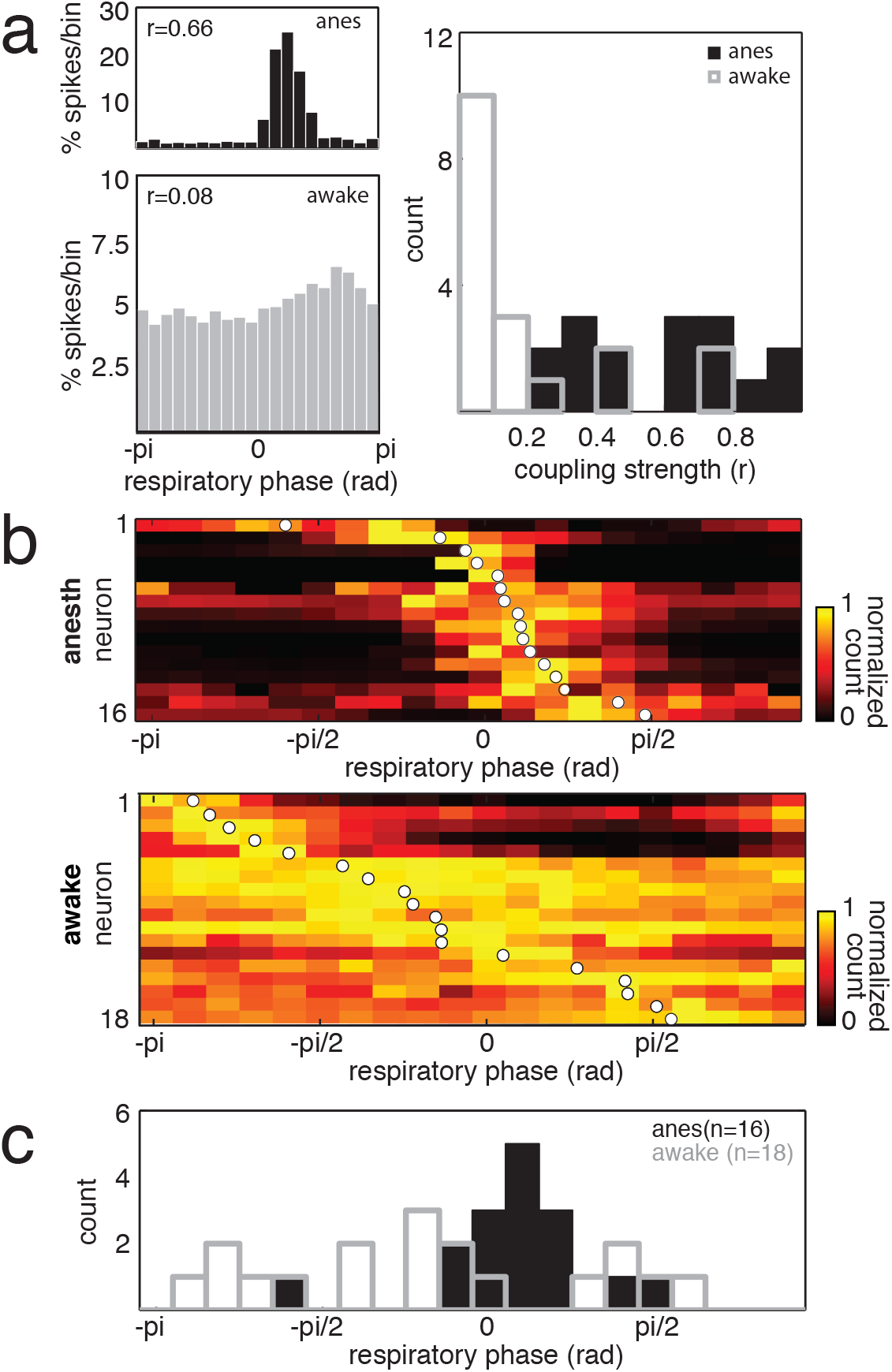
GCs show strong synchronous coupling to respiration in anesthesia and weak divergent coupling to respiration in wakefulness. (a) Respiratory phase coupling histograms for a representative (near mean coupling strength) anesthetized state GC (black) and an awake state GC (gray). Histograms show the probability of spikes occurring in each pi/10 bin of the respiratory cycle. Zero corresponds to the peak of inspiration. The right panel shows a histogram (bin size = 0.1) of respiratory coupling strength (*r*) for anesthetized state GCs (black) and awake state GCs (gray). GCs in the anesthetized state are significantly more coupled to breathing than cells in the awake state (*p* < 0.001, U test). (b) Heatmap depicting baseline respiratory phase spike histograms for all anesthetized state GCs (top panel) and all awake state GCs (lower panel) (bin size = pi/10). Each row is a cell, and for each cell, spike count is normalized to the maximal bin for that cell. White dots denote the mean phase angle of firing for each cell. (c) Histogram of mean firing phase for all cells under anesthesia (black) and during wakefulness (gray). The two distributions were significantly different (*p* < 0.01, KS test).

In addition to being strongly coupled to the breathing rhythm, GCs during anesthesia were as a population synchronized to a narrow window of the respiratory phase corresponding to the period just after the peak of inspiration (Fig. 3b, top). Interestingly, this synchrony was not evident in the population of GCs recorded during wakefulness. Instead, in the awake state GCs displayed divergence in their mean coupling phase angles, covering a much broader range of the respiratory cycle. The distributions of mean phase angles were significantly different for awake state GCs and anesthetized state GCs (Fig. 3c) (*p* < 0.01, Kolmogorov Smirnov test).

Mitral/tufted cells are phase coupled at baseline and frequently shift their coupling phase angle in the presence of an odor stimulus. The direction and magnitude of the phase angle shift depends on odor stimulus identity^11, 42–44^. We therefore assessed whether phase coupling angle in GCs was stimulus-dependent in awake or anesthetized mice. We first selected neurons that showed significant respiratory coupling at baseline (16/16 anesthetized state GCs and 16/18 awake state GCs). For each baseline coupled cell, we identified all odors for which spiking during the trials of that odor was significantly coupled to respiration. This yielded 300 cell-odor pairs (from 15 cells) in anesthetized state GCs and 45 cell-odor pairs (from 10 cells) in awake state GCs. Fig. 4a shows data from an example anesthetized state GC and an example awake state GC, comparing the baseline activity phase histogram with the stimulus-specific phase histograms for all odors that evoked significantly phase coupled activity from that cell. In each case, the stimulus-specific coupling phases for all odors clustered tightly around the mean baseline coupling phase. We quantified this close correspondence between baseline and stimulus-driven coupling by correlating the phase histograms of each cell-odor pair with the baseline phase histogram of its parent cell. A large majority of cell-odor pairs showed a significant positive correlation between their stimulus-driven histogram and baseline phase histogram (276/300 cell-odor pairs from anesthetized state GCs and 32/45 cell-odor pairs from awake state GCs), and the mean cell-odor pair correlation for both populations was high (anesthesia corr. coeff. r: 0.77 ± 0.2; awake corr. coeff. r: 0.57 ± 0.2).

**Figure 4.**
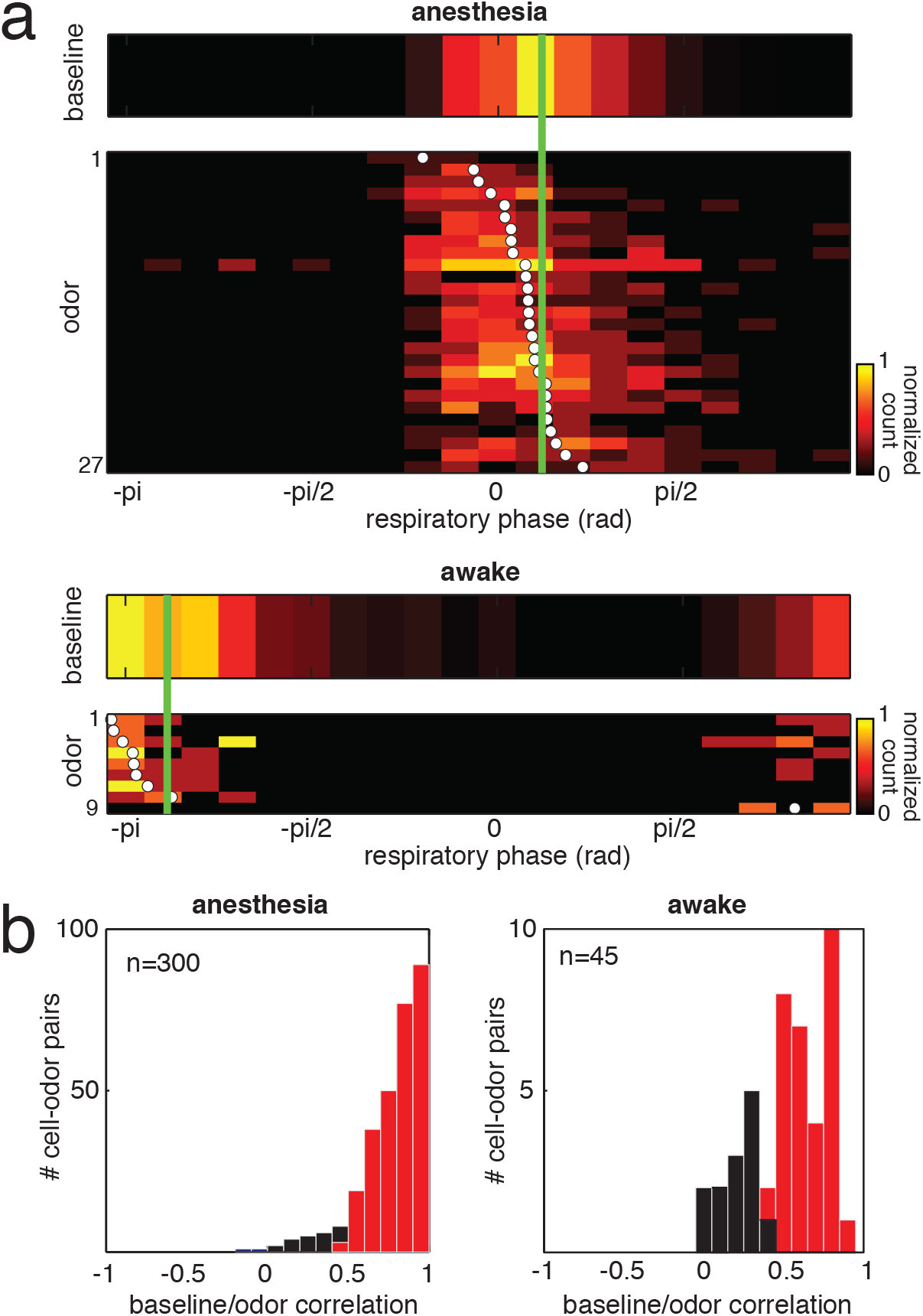
Respiratory coupling phase of GC spiking is unaffected by odor stimuli. (a) Heatmaps depicting baseline and odor-selective respiratory phase spike histograms for an example anesthetized state GC (top panels) and an example awake state GC (lower panels) (bin size = pi/10). Each row is an odor, and for each odor, spike count is normalized to the maximal bin for that odor (or baseline). Only cell-odor pairs for which there was significant coupling in both the baseline and odor data are plotted. White dots denote the mean phase angle of firing for each odor and the green line denotes the mean baseline firing phase for each cell. (b) Correlation of baseline and odor phase histograms for all cell-odor pairs revealed that the great majority of cell-odor pairs were highly and significantly correlated at baseline and during the odor (red). This was true in both anesthesia and wakefulness, and demonstrates that unlike MTs, GCs do not encode odor identity as shifts in phase of respiratory coupling.

Thus contrary to our prediction, GCs are not strongly coupled to respiration during wakefulness. Taken together with our further observation that individual GCs have weak and diverse phase relationships with breathing and that this relationship is insensitive to odors, we conclude it is unlikely that GCs play a dominant role in influencing respiratory locking in MTs during wakefulness. Moreover, GCs do not appear to even merely follow the respiratory locking of MTs.

## Odor responses in GCs are independent of respiration during wakefulness

The analysis we describe above suggests that odor information carried by GCs is independent of the breath cycle during wakefulness. If so, that argues against a role for GCs in patterning MT firing according to respiration. We therefore compared the dependence of odor coding in the two behavior states on respiratory phase by syncing GC spiking data to breathing.

Fig. 5a shows 2-D histograms of the response of the same anesthetized state GC analyzed with respect to time relative to stimulus onset or with respect to breath cycle number relative to the first inhalation after stimulus onset. Averaging firing rates in time bins relative to stimulus onset (Fig. 5a, left histogram) revealed little structure in the response to odors, as in Fig. 2a. The tuning curve computed from these data during the stimulus was very flat (Fig. 5a, left tuning curve). Constructing the histogram such that each bin is a breath, with breath 1 corresponding to the first inhalation of the stimulus (see Online Methods), sufficiently aligned the data to reveal clearer and more widespread changes in spike rate per breath relative to the baseline per breath spiking rate (Fig. 5a, middle histogram). However, the tuning curve computed from these data was still quite flat (Fig. 5a, middle tuning curve).

**Figure 5.**
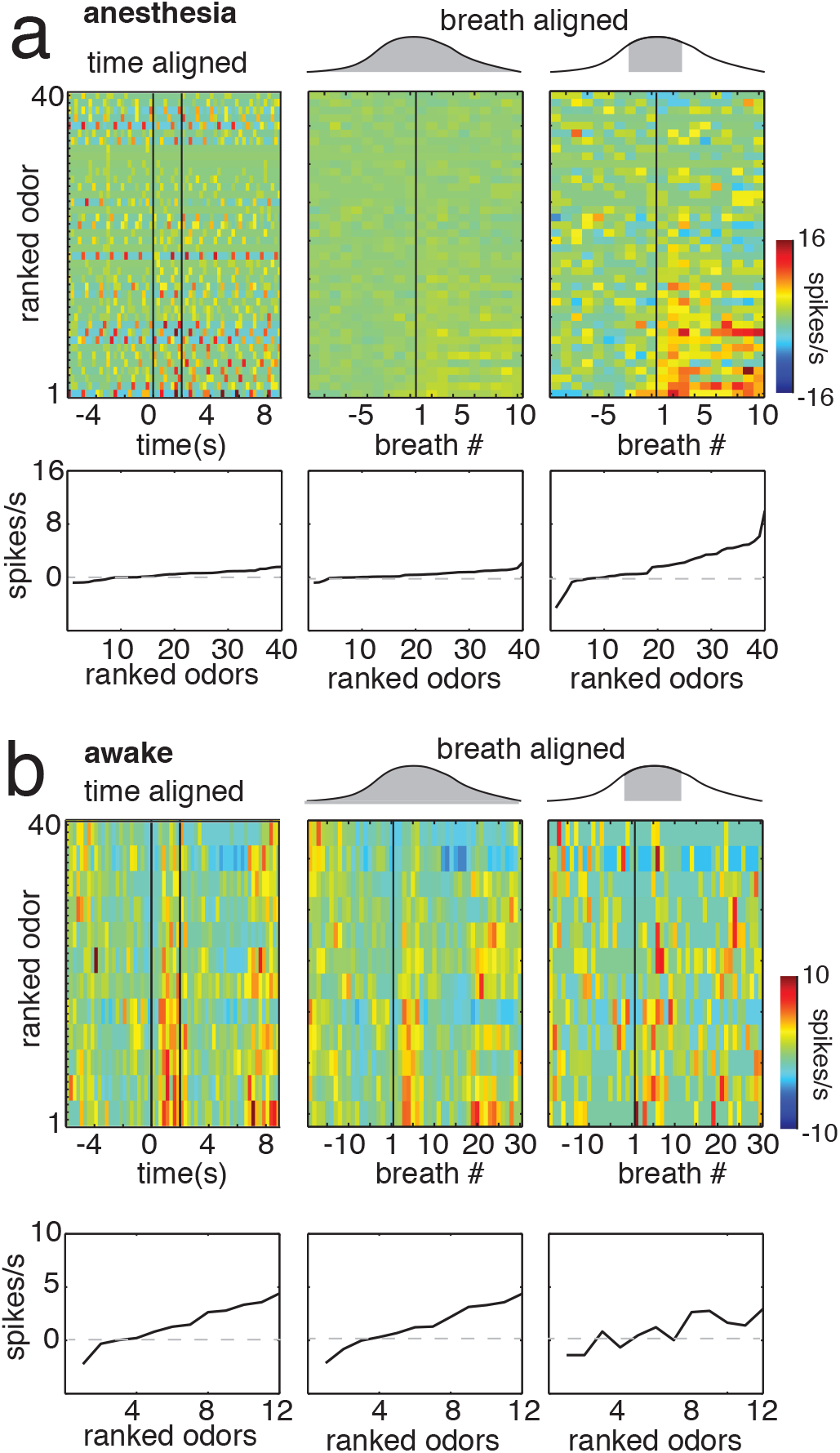
Temporal structure of odor responses differs in anesthetized state and awake state GCs. (a) PSTHs from a representative anesthetized state GC computed with three different methods. The left PSTH depicts mean firing in 250 ms time bins relative to the stimulus onset. The middle PSTH depicts mean firing over each entire breath relative to the first breath after stimulus onset. ‘Full breath’ refers to data calculated in this fashion. The right PSTH is similarly computed, but is restricted to firing in a discrete window of each breath (25% of the respiratory cycle). ‘Partial breath’ refers to data calculated in this fashion. Structure in the cell’s response to stimuli becomes more apparent after syncing to the breath cycle and especially after restricting analysis to part of the breath. Below each PSTH is plotted the corresponding ranked tuning curve computed with each method. Tuning curves and PSTHs are ranked according to strength of the response during the odor. (b) Corresponding data from a representative awake state GC.

Given our observation that most spiking in anesthetized state GCs occurs in a narrow window of the breath cycle, we assessed whether considering only part of the breath cycle improved stimulus discrimination, and whether the quality of stimulus discrimination was a function of which part of the breath is considered. Restricting our firing rate computation to 25% of the breath cycle revealed widespread and robust changes in firing rate per breath to many odors (Fig. 5a, right histogram). This difference was reflected by a much steeper tuning curve (Fig. 5a, right tuning curve). The degree of benefit from this analysis was a function of which 25% of the breath cycle was analyzed. Sliding the 25% window over the breath cycle revealed a sharp peak in tuning curve variance (Fig. 5a, right panel, blue trace) that corresponded with a sharp peak in correlation between the partial breath tuning curve and the time aligned tuning curve (Fig. 5a, right panel, red trace).

Visual inspection of the three histograms produced in the same manner from an example awake state GC suggested that the quality of stimulus discrimination did not benefit from analysis of firing rate changes during a restricted respiratory phase (Fig. 5b). Additionally, high correlation between the partial breath tuning curve and the time-aligned tuning curve was maintained throughout most of the respiratory cycle. These qualitative observations suggest that odor information encoded by GCs is distributed throughout the respiratory cycle during wakefulness but is concentrated in a narrow window during anesthesia.

To quantify this difference for all cells, we measured the dependence of partial breath tuning curve variance and correlation on respiratory phase angle by measuring the coupling of these variables to the breath cycle. Both measures tended to show peaks at specific points in the breath cycle under anesthesia, while during wakefulness both measures tended to be invariant with respiration (Fig. 6a). As a population, compared to anesthetized GCs, awake state GCs exhibited significantly lower respiratory coupling of tuning curve variance (anesthesia coupling strength: 0.59 ± 0.2, n = 15; awake coupling strength: 0.29 ± 0.1, n = 16; *p* < 0.001, t test) and correlation to time aligned tuning curves (anesthesia coupling strength: 0.49 ± 0.2, n = 15; awake coupling strength: 0.29 ± 0.1, n = 16; *p* < 0.01, t test) (Fig. 6b and c).

**Figure 6.**
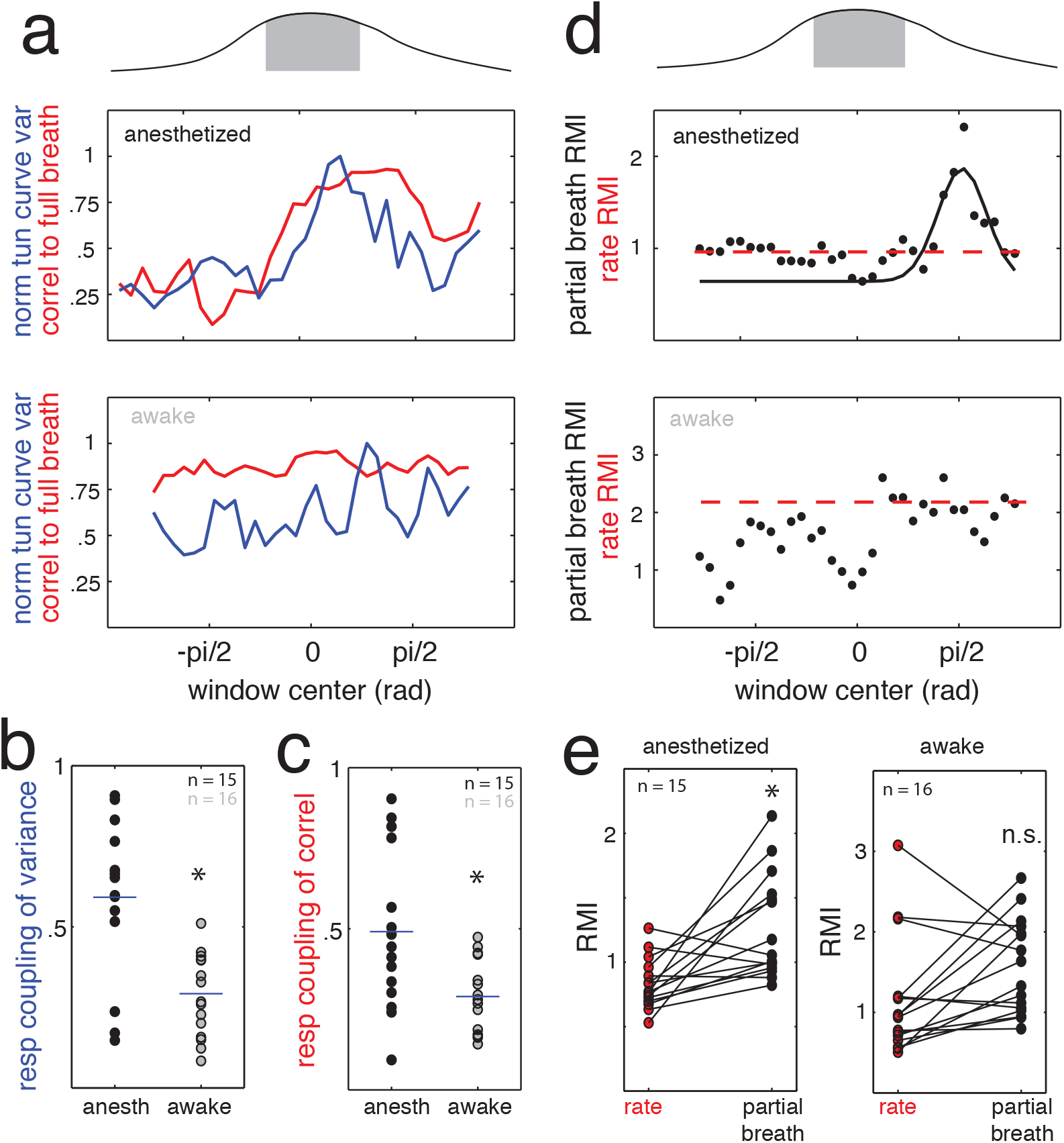
Information about odors depends on the respiratory cycle in anesthetized state GCs but not in awake state GCs. (a) Partial breath tuning curve variance (blue) and correlation to the full breath tuning curve (red) are plotted as a function of respiratory phase for a representative (near mean) anesthetized state GC (top panel) and an awake state GC (lower panel). For anesthetized state GCs, these measures of tuning curve robustness and accuracy were coupled to respiratory phase, while they were independent of the breath cycle in awake state GCs. (b,c) Scatterplots comparing the strength of coupling between partial breath tuning curve variance (b) and correlation to the full breath tuning curve (c) for anesthetized state GCs and awake state GCs. Awake state GCs showed significantly weaker coupling than anesthetized state GCs for both measures (*p* < 0.05, t test). (d) RMI values for partial breath tuning curves are plotted as a function of respiratory phase for a representative (near mean) anesthetized state GC (top panel) and an awake state GC (lower panel). Where possible, the data were fit with a Gaussian (black curve) to identify the peak value as the maximal ‘partial breath RMI’. If the data could not be fit with a Gaussian, the mean value was used. The dashed red line indicates the RMI value computed from firing rate without respect to breathing or ‘rate RMI’. (e) Scatterplots comparing the rate RMI values and the partial breath RMI values for anesthetized state GCs (left) and awake state GCs (right). Anesthetized state GCs showed significantly increased RMI values computed from partial breath tuning curves (*p* < 0.05, one way ANOVA comparing rate, full breath, and partial breath RMI values). Awake state GCs showed no significant difference in this comparison.

We further quantified whether partial breath tuning curves improved stimulus discrimination by measuring response modulation index (RMI) as a function of respiratory phase angle. We computed RMI values for time aligned data as well as data in a sliding window restricted to 25% of the breath cycle (see Online Methods). Where possible, partial breath RMI values were fit with a Gaussian function, and the peak of the Gaussian was taken as the peak RMI for partial breath analysis (e.g. Fig. 6d, top). For cells that could not be well fit with a Gaussian function (e.g. Fig. 6d, bottom), the mean RMI value over the respiratory cycle was used. Considered as a population, anesthetized state GCs exhibited significantly higher RMI values for partial breath as compared to time-aligned analysis (time-aligned firing rate RMI: 0.83 ± 0.2; partial breath-aligned firing rate RMI: 1.27 ± 0.4; n = 15, *p* < 0.05, ANOVA), whereas awake state GCs did not (time-aligned firing rate RMI: 1.56 ± 0.6; partial breath-aligned firing rate RMI: 1.27 ± 0.4; n = 15, *p* > 0.05, ANOVA).

These data indicate that odorant information is distributed over the entire respiratory cycle during wakefulness. Likewise, ongoing activity in awake state GCs has limited temporal structure. As a result, odor responses and ongoing activity exhibit slower and steadier dynamics that do not reflect a strong influence of respiration. We therefore find no evidence that GCs contribute to any temporal patterns in MTs related to breathing.

## Discussion

Here we investigated state-dependent activity in GCs of the main olfactory bulb. Granule cells are the target of extensive central feedback projections and make reciprocal dendrodendritic synapses onto mitral/tufted cells (MTs). They are therefore well poised to be critical effectors of state modulation of MOB output. Indeed, we found dramatic differences in ongoing and stimulus-driven activity between GCs recorded in anesthetized and awake mice. These differences likely have important consequences for odor coding. Specifically, recordings during wakefulness suggest that GCs are likely to have more extensive lateral interactions with mitral cells, but are unlikely to strongly shape their temporal dynamics.

The activity of GCs under anesthesia was characterized by temporally sparse bursts that were synchronized across cells. As a result, spiking and therefore odorant information is concentrated in a narrow window aligned with respiration. Given the limited sampling rate of laser scanning microscopy, this temporal sparseness may explain why odor responses were not readily observed from GCs of anesthetized mice in a recent two photon imaging study^16^. This is also relatively consistent with whole-cell recordings from GCs under anesthesia, which reveal a highly nonlinear spike threshold mechanism that enforces temporal sparseness despite strong subthreshold input^28^. However, our data do stand in contrast with the large, sustained changes in firing rate reported for GCs in one recent study of anesthetized mice^29^. Perhaps this is related to selection of an anesthetic, however we observed similar activity with two common anesthesia regimes (Supplemental Fig. 6).

The awake state was characterized by two major features. First, GCs exhibited higher firing rates and stronger, more widespread odor responses. Based on the fact that GCs define a circuit enabling mutual inhibition between neighboring MTs, it is widely held that they tune responses of individual MTs and sparsen population representations of odors^16, 21, 26, 27^. The observation that GCs respond to more odors and more sensitively discriminate among them implies that under wakefulness they play a greater role in sculpting MT selectivity through augmented lateral interactions. This conclusion is broadly consistent with experimental and theoretical observations regarding MT activity in awake animals and GC function^8, 9, 16, 26, 45–47^.

Second, the awake state was characterized by reduced coupling to the breathing rhythm and divergence of respiratory coupling phase across cells. Consequently, unlike under anesthesia, information regarding odorant identity was distributed throughout the breathing cycle. The observation that GCs largely fire with little regard for the respiratory cycle leads us to conclude that it is unlikely they play a significant role in the enforcement of the respiratory locking exhibited by MTs in awake animals.

When compared with the temporal structure of activity observed in MTs in awake animals, our results are surprising in two respects. First, several studies have reported reliable stimulus-specific shifts in respiratory phase coupling of MTs^11, 42–44^. Granule cells have naturally been proposed as a likely contributor to this effect^42, 43^. However, in our data, GCs have a characteristic coupling phase that is invariant to stimuli in both the awake and anesthetized states. Thus, if odor-specific phase shifts are accomplished via GC-mediated feedback inhibition, it is more likely to arise from modulation of the amplitude (as opposed to timing) of inhibitory pulses of specific GC ensembles to push and pull the phase of MT coupling. Second, the weak respiratory coupling and apparent slow rate coding we observed in GCs during wakefulness stands in contrast to the high temporal precision of MTs recorded under similar conditions^11, 12^. Because of the strength of reciprocal coupling between GC and MTs, one might expect that GCs sculpt or at least reflect the precise MT bursts that tile the respiratory cycle in awake rodents^12^. Nonetheless, we failed to find evidence that GCs participate substantially in this phase locking or even passively follow breathing. Although we cannot completely exclude highly cooperative population mechanisms for creating temporal structure GCs, our observations limit the potential influence of one or a small number of GCs on respiratory patterning in MTs.

Population activity of MTs is read out by the olfactory cortex. Considerable evidence argues that the cortex spatially integrates input from multiple glomeruli ^48–54^, but the temporal integration properties of the olfactory cortex are less clear and are under active investigation^55, 56^. Synchronous inhibition seems likely to entrain synchronous MT output which may be best extracted by coincidence detection, whereas the slower firing rate changes seen in our awake data may be best served by temporal integration upon readout. Potential support for regulated temporal integration comes from recent evidence revealing the existence of an intriguing mechanism for modulating MT synchrony to encode stimulus associations^15^.

Finally, while the state-dependent modulation we observed was between anesthesia and passive wakefulness, our results demonstrate remarkable lability of the MOB interneuron network. This capacity for flexibility is therefore likely exerted to optimize neural processing for a wide range of behavioral conditions.

## Acknowledgements

The authors wish to thank A. Kepecs, S. Ranade, and J. Sanders for technical advice, and A. Zador, B. Li, R. Mooney, A. Fontanini, R. Froemke, and Y. Ben-Shaul for comments on an earlier version of the manuscript.

## Author Contributions

BNC and BYL contributed equally. SDS supervised the project and designed the experiments together with BNC and BYL. All authors developed the methods and participated in collecting the data. SDS, BNC, and BYL analyzed the data and wrote the paper.

## METHODS

### Animals

Experiments were performed on adult (aged 6–12 weeks) male C57Bl/6 mice (Jackson Laboratory). Mice were maintained on a 12/12 h light dark cycle (lights on 0700 h) and received food ad libitum. For awake recording, water bottles were removed from the cage 24 h prior to each session, and animals only had access to water through the recording apparatus lick port (Supplementary Figure 1). All procedures were conducted in accordance with the National Institutes of Health’s *Guide for the Care and Use of Laboratory Animals* and approved by the Cold Spring Harbor Laboratory Institutional Animal Care and Use Committee.

### Surgery and Measurement of Respiration

For awake recordings, animals were anesthetized with an 80:20 mixture (1.25 ml/kg) of ketamine (100 mg/ml) and xylazine (20 mg/ml) and stabilized in a stereotaxic frame. A head bar was affixed to the skull immediately posterior to the coronal suture using adhesive luting cement (Parkell, Inc) and methyl methacrylate-based dental cement (TEETS). For additional support, four machine screws (Amazon Supply) were secured to the skull prior to application of the luting cement. To assist in the measurement of breathing, a cannula made from polyimide tubing (ID 0.0319, Amazon Supply) was unilaterally implanted in the nasal cavity rostral to the olfactory epithelium using dental cement. When not recording, this cannula was plugged with a stainless steel dummy insert. In awake animals, breathing was measured using Teflon insulated thermocouples (0.13 mm OMEGA, part # 5TC-TT-J-36-36) placed acutely in the cannula, and data were amplified (Brownlee 410), digitally filtered, and acquired at 1 kHz using Spike2 (CED).

Some recordings were conducted in animals that remained continually anesthetized. For these recordings, animals were anesthetized using either ketamine/xylazine or isoflurane (∼1% in pure O_2_). Breathing was recorded using a foil strain gauge (Omega Engineering) placed on the animal’s trunk.

### Odor Stimuli

Odor stimuli were delivered as described (Shea et al, 2008). Stimuli consisted of a number of different monomolecular odors as well as natural food odorants (Supplementary Table 1) diluted to 1% V/V in mineral oil. An additional 1:10 flow dilution in our olfactometer resulted in a final concentration at the nose of 0.1% saturated vapor. Typical odor presentations in the awake state consisted of 3–4 repetitions of 7–11 different odors. In anesthetized animals, the stability of recording permitted the presentation of many more odors (20–40). All odors were presented for a 2 s duration followed by a 13 s interstimulus interval (ISI). In order to keep awake animals engaged and comfortable, water (20 ul) was delivered through a water port at the end of each odor presentation. For assessment of cell firing between awake and anesthetized conditions in the same animal, isoflurane anesthetic (∼1%) was delivered to the animal through the odor port in addition to the odors.

### Electrophysiology

Mice were anesthetized (1% isoflurane), and the MOB was exposed with a small craniotomy. Awake mice were head fixed via the attached head bar over a Styrofoam ball that permitted them to walk and run in one dimension (backwards/forwards). The ball was suspended above the air table and attached to a rotary encoder (Sparkfun) for measurement of locomotor activity. In vivo, blind loose-patch recordings were conducted using borosilicate micropipettes (15–25 MΩ) tip-filled with intracellular solution (125 mM potassium gluconate, 10 mM potassium chloride, 2 mM magnesium chloride, 10 mM HEPES, pH 7.2) and further filled with 1.5% neurobiotin. Neuronal spiking was recorded using a BA-03X bridge amplifier (npi Electronic Instruments), low pass filtered at 3 kHz, and digitized at 10 kHz. Data were acquired using Spike2 software and analyzed offline using Spike2 and Matlab. Following each recording, cells were filled using positive current injection (+700 pA; 1 Hz) of neurobiotin for 15–25 min.

**Table 1.**
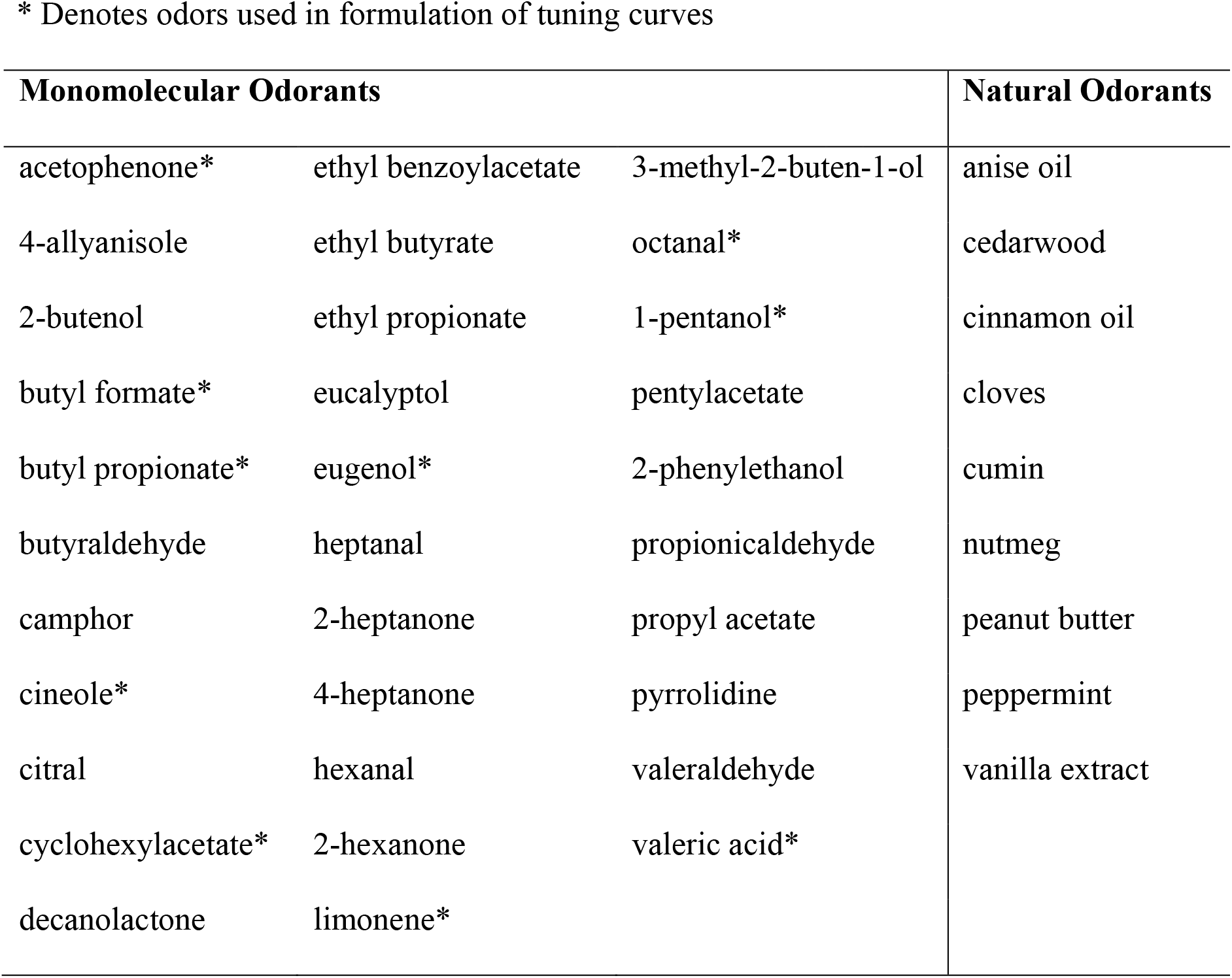
List of Odorants

\* Denotes odors used in formulation of tuning curves
| Monomolecular Odorants |  |  | Natural Odorants |
| --- | --- | --- | --- |
| acetophenone* | ethyl benzoylacetate | 3-methyl-2-buten-1-ol | anise oil |
| 4-allyl anisole | ethyl butyrate | octanal* | cedarwood |
| 2-butenol | ethyl propionate | 1-pentanol* | cinnamon oil |
| butyl formate* | eucalyptol | pentylacetate | cloves |
| butyl propionate* | eugenol* | 2-phenylethanol | cumin |
| butyraldehyde | heptanal | propionaldehyde | nutmeg |
| camphor | 2-heptanone | propyl acetate | peanut butter |
| cineole* | 4-heptanone | pyrrolidine | peppermint |
| citral | hexanal | valeraldehyde | vanilla extract |
| cyclohexylacetate* | 2-hexanone | valeric acid* |  |
| decanolactone | limonene* |  |  |

### Immunohistochemistry

Following recording, animals were sacrificed using an overdose of sodium pentobarbital (Euthasol) and transcardially perfused with PBS followed by 4% paraformaldehyde. Brains were extracted in the skull and stored in paraformaldehyde for ∼12−16 h followed by 30% sucrose for ∼ 24 h. Olfactory bulb sections (100 um) were treated according to standard immunohistochemistry procedures using 1:667 streptavidin Alexa 594 (Invitrogen). Sections were first viewed under epifluorescence (Olympus BX43), and images were further taken using a LSM 710 laser scanning confocal microscope (Carl Zeiss). Only cells identified as granule cells or as being in the granule cell layer (Supplementary Figure 3) are reported here.

### Data Analysis

Recordings sometimes included contamination from other spikes or artifacts, so manual spike sorting with Spike2 (CED) was used to isolate single unit spike trains. All subsequent analyses were performed in Matlab (Mathworks). Mean ongoing firing rate were calculated from the 6 s period just prior to each stimulus. This baseline was subtracted from the spike rates measured during the 2 s stimulus and during the 1 s after the stimulus offset to compute response strength by firing rate.

Statistical significance of responses to individual odors was assessed with a bootstrap procedure as follows. If *n* trials were collected with the response window length *t*, then a distribution was created by sampling *n* length *t* windows from the full spike record 100000 times and taking the mean deviation of each window from the spike rate measured in the prior 6 s. Responses that were in the top or bottom 2.5% of this distribution were deemed significantly excitatory or inhibitory respectively. Our response modulation index (RMI) was computed as in reference 34, with the exception that in our data, responses could be negative, thus no correction for non-normally distributed data was required. Briefly, RMI for each cell was calculated as the ratio of the variance across stimuli over the variance across all trials:

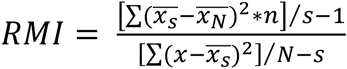

where *x* is the response on one trial of an odor, *n* is the number of trials for one odor, *N* is the total number of trials across all odors, *s* is the number of stimuli, 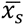 is the mean response to an individual stimulus, and 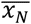 is the mean response for all trials across all odors. The strength and phase of coupling between spiking and respiration was computed using the publicly available Matlab library Circstat (ref. 41). Statistical significance of coupling for a given stretch of data was determined with a Rayleigh test (ref. 41).

Breath synchronized odor responses were analyzed as follows. The time of each inspiratory peak in the entire record for a given cell was identified. Then, spike rate was computed for each breath in a window between the halfway point to the prior breath and the halfway point to the subsequent breath. The first inspiration to peak after the onset of the odor was designated as breath 1. Baseline per breath firing was designated as the mean per breath firing rate for the 20 breaths immediately prior to the odor trial. This baseline value was subtracted from the mean per breath firing rate for the breaths that occurred during the stimulus to compute breath synchronized response strength. “Whole breath” tuning curves and histograms were constructed in this way. “Partial breath” tuning curves and histograms were constructed in the same manner but considering only a specific window of 25% of the breath cycle. Cycle-dependent structure in the tuning curves and histograms was assessed by sliding this window over a range of positions on the breath cycle.

---

**Supplementary Figure 1.**
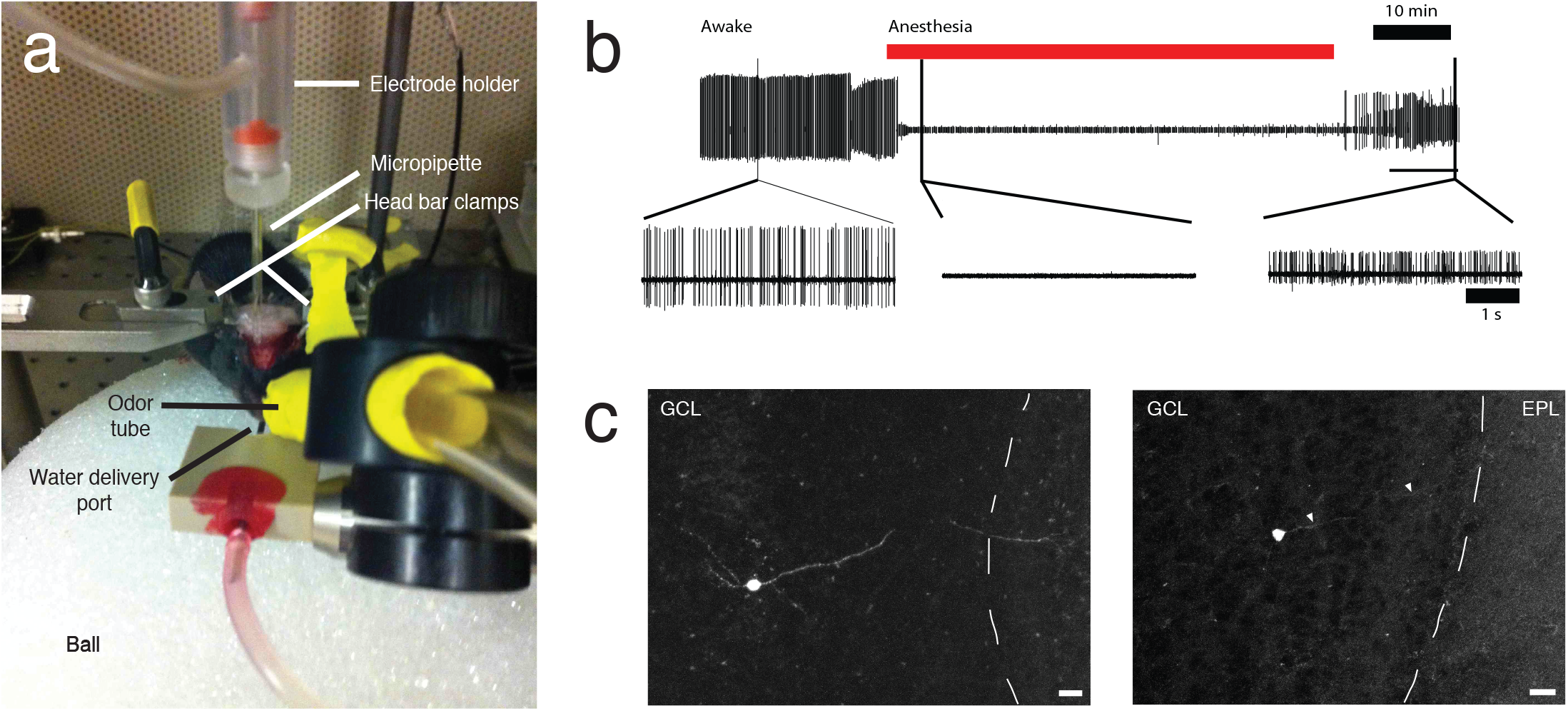
Physiology and anatomical recovery methods for GCs in awake mice (a) Mice were head-fixed on a Styrofoam ball by clamping a head bar affixed to the skull with dental cement. Mice could move freely in 1 dimension (forward or backward). An odor delivery tube (yellow) attached to an olfactometer was placed ∼ 5 cm from the animal’s nose. Both odor and isoflurane (during anesthesia) were delivered through this tube. Water reward was delivered through a lick-port positioned below the odor delivery tube. (b) A typical recording from a GC across awake and anesthetized states. Upon several presentations of odor stimuli, animals were transitioned from awake to asleep using isoflurane anesthetic (red line). Where recording stability allowed, recordings were then conducted from the same cell in the anesthetized animal for several odor presentations. Then, anesthetic was turned off, and recording continued as the animal recovered. Timing of odor presentations are not shown. (c) Granule cells in the awake animal were labeled using the same methods as for granule cells in the anesthetized animal. Movement limited cell fill times, but anatomical location, cell size, and the presence of an EPL projecting apical dendrite (arrows) confirmed GC identity. Scale bar = 20 um. (GCL = granule cell layer, EPL = external plexiform layer).

**Supplementary Figure 2.**
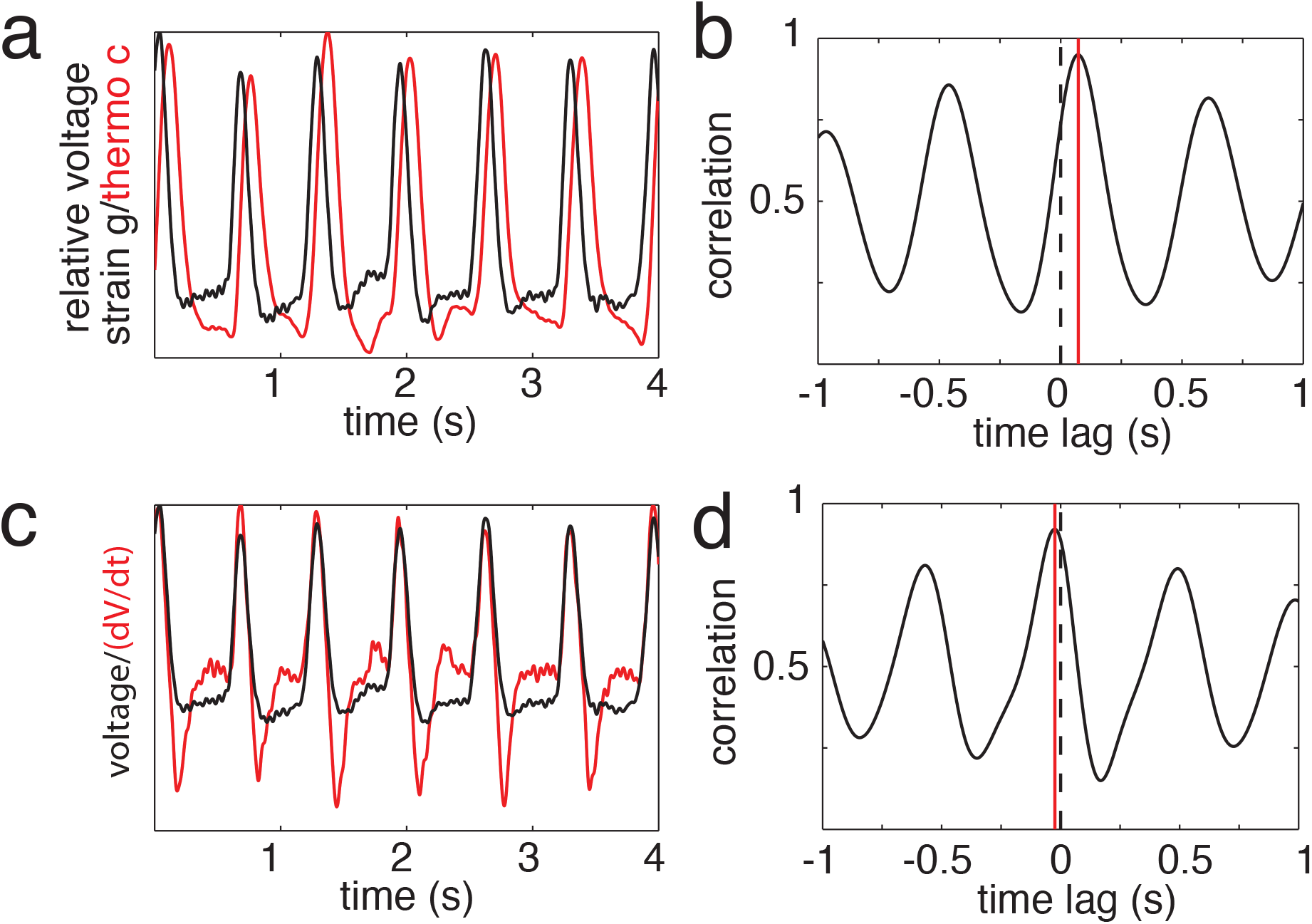
Comparison of two methods for measuring respiration. (a) Example raw traces from simultaneous respiratory recordings using a nasally implanted thermocouple (red) and a foil strain gauge (black). We observed that the peak of the strain gauge signal closely matched the point of maximum rate of change for the thermocouple signal. (b) Cross correlation of the two signals shows a lag of 68 ms for the strain gauge relative to the thermocouple. (c) Comparison of the strain gauge with the differentiated thermocouple signal shows a good match of the signal peaks. (d) Cross correlation of the two signals in (d) shows a greatly reduced lag of 15 ms for the strain gauge relative to the thermocouple. Mean and SD time between peaks was 0 +/− 7 ms.

**Supplementary Figure 3.**
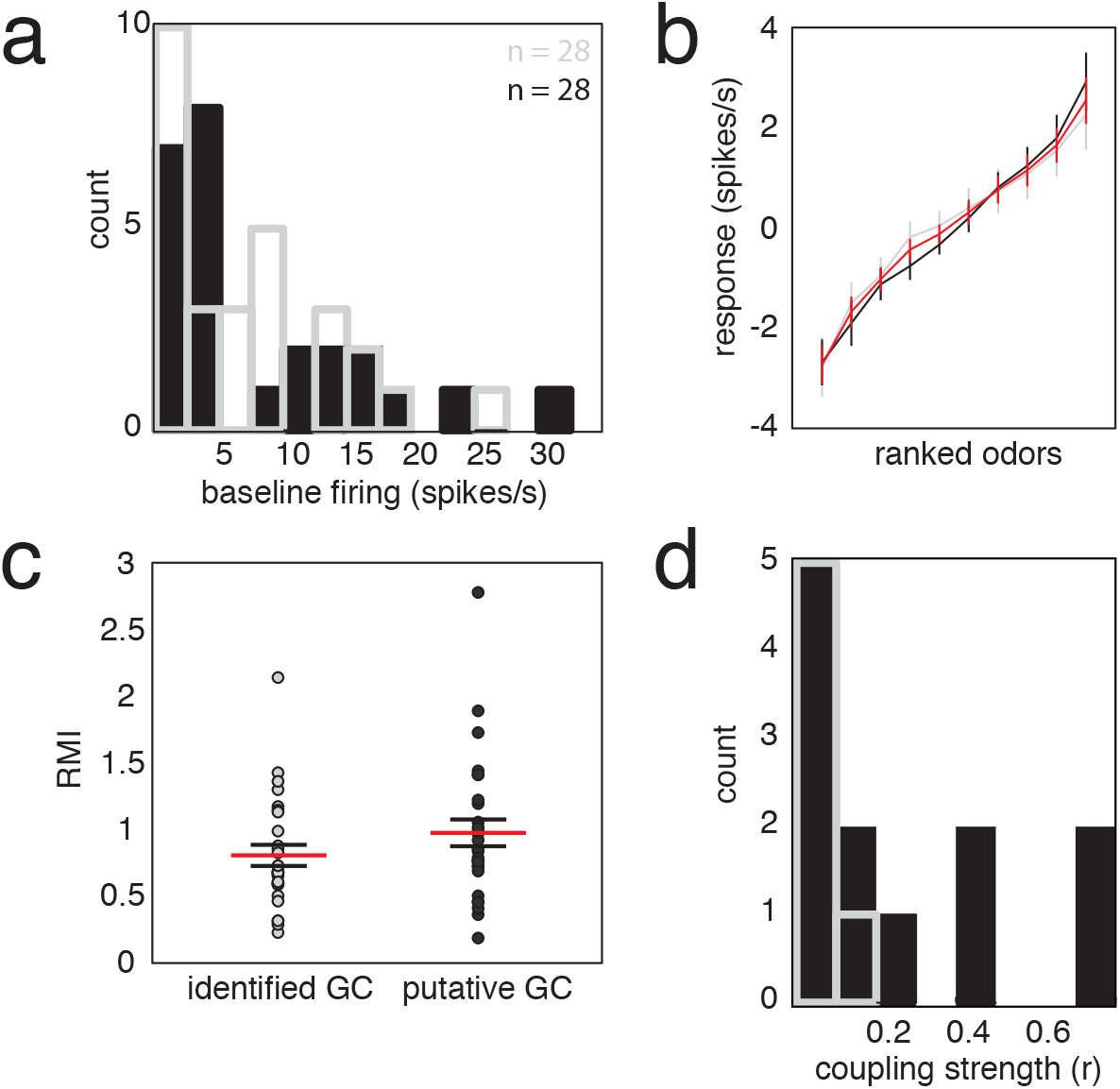
Granule cell layer neurons (putative GCs) and identified GCs do not differ in their properties (a) Histogram of identified GCs (grey) and putative GCs (black) shows no difference in spontaneous activity between the two cell populations. Identified GCs had a mean baseline rate 7.30 ± 1.3 spikes/s while putative GCs had a mean baseline rate of 7.99 ± 1.5. (b) Ranked tuning curves for identified and putative GCs are similar. Identified GC curve is plotted in grey, putative GCs are plotted in black, and the combined curve is plotted in red. Only spike rates during the odor presentation are shown. (c) Response modulation index (RMI, see Online Methods) does not differ between the two groups. Each data point represents the RMI value for one cell. Mean RMI identified GC = 0.81 ± 0.08; mean RMI putative GC = 0.98 ± 0.10. (d) Coupling of spiking to breathing is not significantly different in the two groups. In both the identified GC and putative GC groups, most cells show little coupling with breathing.

**Supplementary Figure 4.**
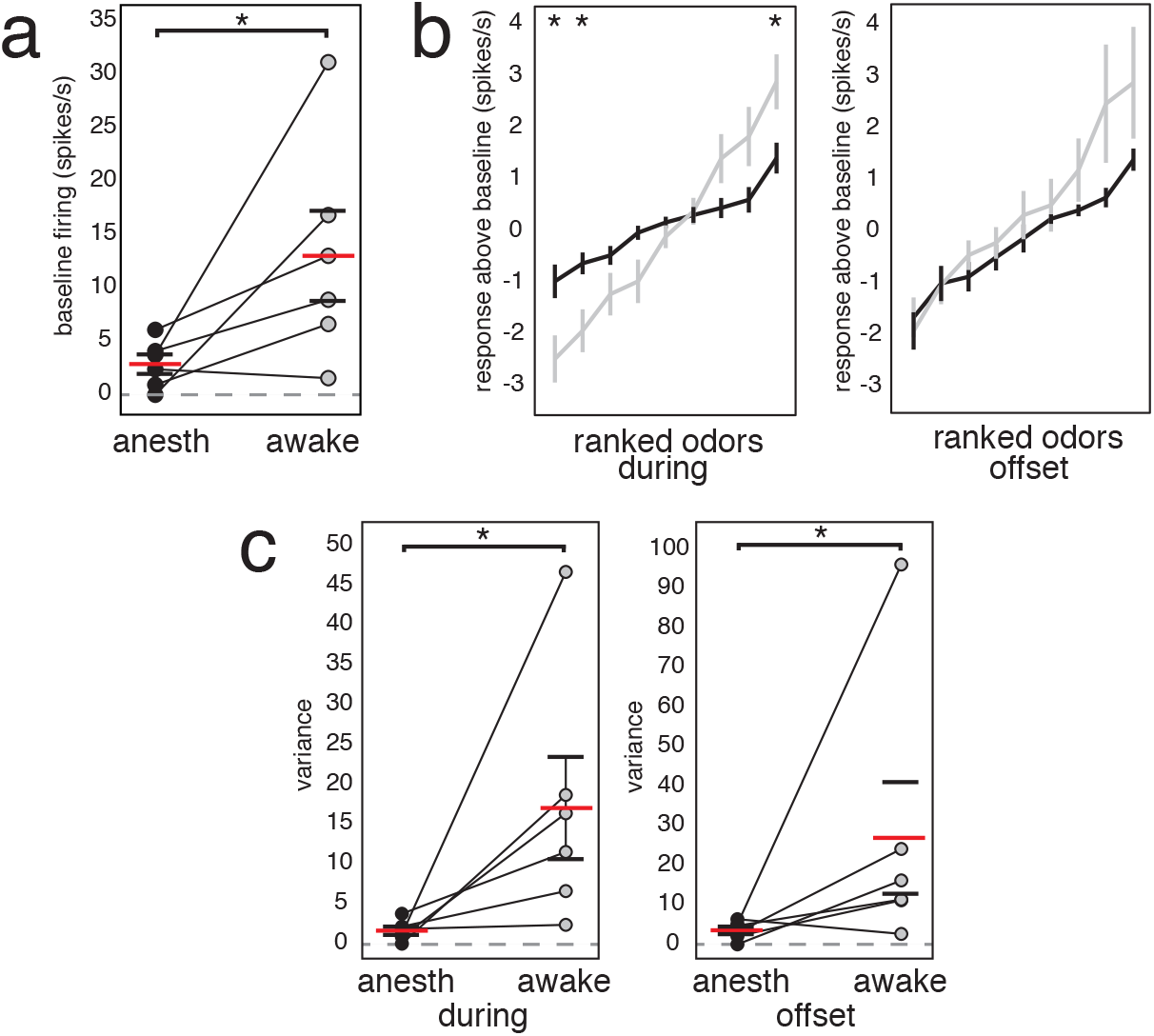
Recordings from GCs across wakefulness state show state-dependent activity changes within individual GCs. In a subpopulation of cells (n = 6), recordings were made from the same cell in both the awake and isoflurane anesthetized animal. (a) Spontaneous activity significantly increased in GCs across the anesthetized (black) to awake (grey) transition (Mann Whitney U test, p = 0.026). Mean firing rate ± SEM (red line and black lines) in the anesthetized state = 2.86 ± 0.91; mean firing rate in the awake state = 13.00 ± 4.21. Paired data are plotted for each cell with the connecting line linking mean firing rate for an individual cell in each state. (b) Ranked tuning curves during 2 s odor presentation and 1 s after odor offset are plotted for transitional cells in both the anesthetized (black line) and awake (grey line) state. (c) Tuning curve variance during both the 2 s odor presentation and 1 s following odor offset was greater in the awake state (during = 17.04 ± 6.42; after = 26.94 ± 14.13) than in the anesthetized state (during = 1.64 ± 0.53; after = 3.53 ± 0.95). Individual data points represent one cell and mean ± SEM are plotted as horizontal red and black lines.

**Supplementary Figure 5.**
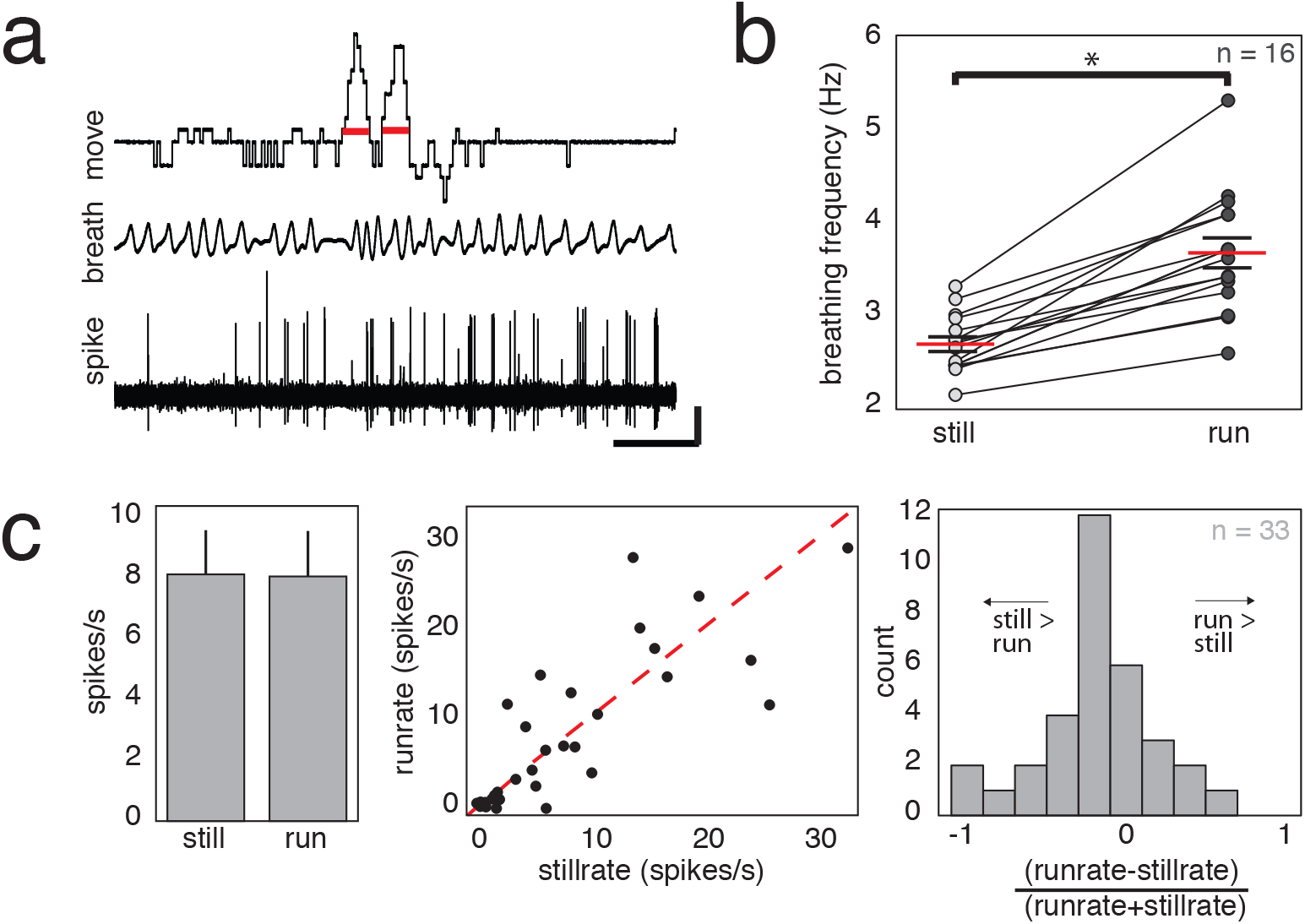
Changes in breathing and GC firing with changes in locomotion. (a) Example trace of GC activity (bottom), respiration (middle), and running velocity (top). Running was measured on an axially rotating Styrofoam ball via a rotary encoder. Locomotor activity was divided into 2 s bins, and running bins with RMS > 2 (red lines; Scale bar is 1 s and 4 cm/s for locomotion, 1 mV for breathing and 1.5 mV for GC spiking. (b) Breathing frequency increases with running. Breathing during the entire recording session was binned into 2 s intervals, and a mean rate during periods of rest or during periods of running was calculated. For all animals, breathing significantly increased during running (Mean breathing rate rest = 2.69 ± 0.08; mean breathing rate during running 3.68 ± 0.16; p < 0.001, Mann-Whitney U test). (c) GC activity is largely invariant with activity. Mean spike rate between periods of rest (8.15 ± 1.46 spikes/s) and periods of increased locomotion (8.08 ± 1.50 spikes/s) was not significantly different (p = 0.71, Mann-Whitney U test; left). Plotting individual data points (middle, left histogram) indicates a sub-population of cells is modulated with activity. Dotted red line runrate = stillrate.

**Supplementary Figure 6.**
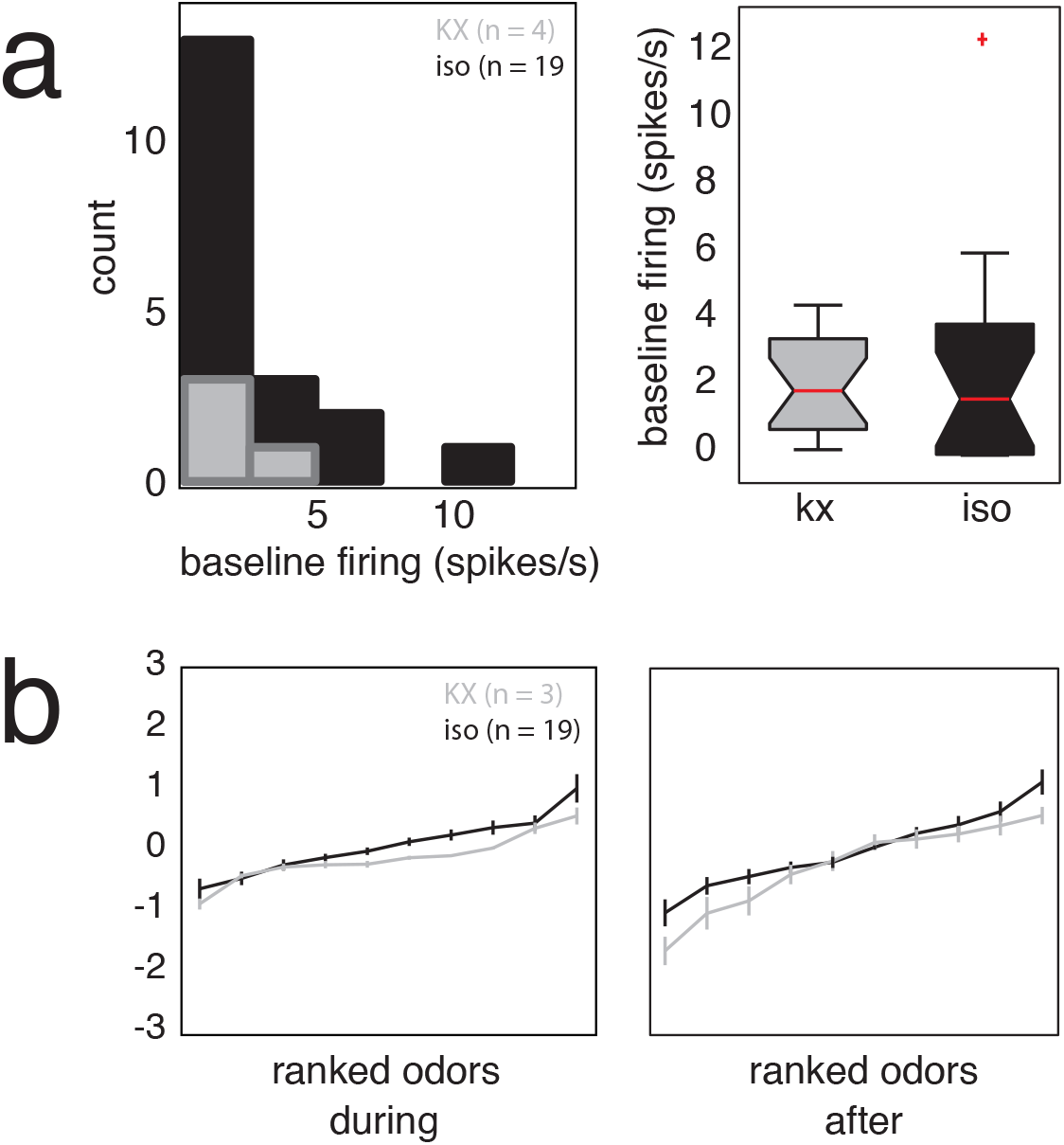
GCs exhibit similar activity under two anesthetic regimes. (a) Histogram of mean spontaneous firing rates from ketamine-xylazine (grey) and isoflurane (black) animals. Mean results are summarized in boxplots (kx = 2.13 ± 0.92, iso = 2.42 ± 0.71). Rates between these two groups did not significantly differ (Mann Whitney U-test, p = 0.65). Kx = ketamine-xylazine, Iso = isoflurane. (b) Ranked tuning curves for both during odor presentation and after odor offset are plotted. Data are represented as the mean ± SEM for each odor and plotted in order from the odor eliciting the most inhibitory response to the odor eliciting the most excitatory response. Curve slope was calculated for both the kx and isoflurane conditions and did not differ across the two anesthetics used (kx during = 0.18, iso during = 0.1611; kx after = 0.24, iso during = 0.25).

